# Characterizing the properties of bisulfite sequencing data: maximizing power and sensitivity to identify between-group differences in DNA methylation

**DOI:** 10.1101/2021.01.22.427791

**Authors:** Dorothea Seiler Vellame, Isabel Castanho, Aisha Dahir, Jonathan Mill, Eilis Hannon

## Abstract

**Background:** The combination of sodium bisulfite treatment with highly-parallel sequencing is a common method for quantifying DNA methylation across the genome. The power to detect between-group differences in DNA methylation using bisulfite-sequencing approaches is influenced by both experimental (e.g. read depth, missing data and sample size) and biological (e.g. mean level of DNA methylation and difference between groups) parameters. There is, however, no consensus about the optimal thresholds for filtering bisulfite sequencing data with implications for the reproducibility of findings in epigenetic epidemiology.

**Results:** We used a large reduced representation bisulfite sequencing (RRBS) dataset to assess the distribution of read depth across DNA methylation sites and the extent of missing data. To investigate how various study variables influence power to identify DNA methylation differences between groups, we developed a framework for simulating bisulfite sequencing data. As expected, sequencing read depth, group size, and the magnitude of DNA methylation difference between groups all impacted upon statistical power. The influence on power was not dependent on one specific parameter, but reflected the combination of study-specific variables. As a resource to the community, we have developed a tool, POWEREDBiSeq, which utilizes our simulation framework to predict study-specific power for the identification of DNAm differences between groups, taking into account user-defined read depth filtering parameters and the minimum sample size per group.

**Conclusions:** Our data-driven approach highlights the importance of filtering bisulfite-sequencing data by minimum read depth and illustrates how the choice of threshold is influenced by the specific study design and the expected differences between groups being compared. The POWEREDBiSeq tool can help users identify the level of data filtering needed to optimize power and aims to improve the reproducibility of bisulfite sequencing studies.

## Background

Epigenetic processes regulate gene expression via modifications to DNA, histone proteins and chromatin without altering the underlying DNA sequence, and there is increasing interest and understanding of the role that epigenetic variation plays in development and disease [1]. The most extensively studied epigenetic modification is DNA methylation (DNAm), the addition of a methyl group to the fifth carbon position of cytosine that occurs primarily, although not exclusively, in the context of cytosine-guanine (CpG) dinucleotides. Despite being traditionally regarded as a mechanism of transcriptional repression, DNAm is actually associated with both increased and decreased gene expression depending upon the genomic context [2], and also plays a role in other transcriptional functions including alternative splicing and promoter usage [3].

Inter-individual variation in DNAm has been associated with cancer [4], brain disorders [5–8], metabolic phenotypes [9, 10] and autoimmune diseases [11]. A number of high-throughput methods have been developed to quantify genome-wide patterns of DNAm, although these differ with regard to enrichment strategy, quantification accuracy and analytical approach [12]. Many approaches are based on the treatment of genomic DNA with sodium bisulfite, which converts unmethylated cytosines into uracil (and subsequently to thymine after amplification) while methylated cytosines are unaffected. The field of epigenetic epidemiology in human cohorts has been facilitated by the development of cost effective, standardized commercial arrays such as the Illumina EPIC Beadchip [13]. Data generated using this platform is relatively straightforward to process and analyze, with a number of standardized software tools and analytical pipelines [14, 15]. These arrays are only currently commonly available for human samples and are limited to capturing predefined genomic positions making up only ~3% of CpG sites in the human genome [16].

For studies requiring greater coverage of the genome, or for the quantification of DNAm in non-human organisms, it is typical to employ highly parallel short read sequencing of bisulfite-treated DNA libraries. A key step in the analytical pipeline of such data is the mapping or alignment of these short sequences back to the genome of interest, a process that is complicated by the degenerated sequence complexity of bisulfite-treated DNA [17]. As well as the need to determine accurately where in the genome a read originates from, the analysis of bisulfite sequencing data involves distinguishing reads mapping to methylated alleles from those mapping to unmethylated alleles. For each cytosine, the level of DNAm is estimated by quantifying the proportion of methylated (C) to unmethylated (T) cytosines from the sequenced reads overlapping that position. Bisulfite sequencing data provides information about cytosine methylation occurring in three distinct sequence contexts: CpG, CHH or CpH sites.

In this paper, we sought to characterize the properties of bisulfite sequencing data with the goal of exploring the experimental variables that influence statistical power and sensitivity to identify differences in DNA methylation in population-based analyses. We define ‘DNAm sites’ as vectors, such that each DNAm site has a ‘DNAm point’ per sample, which incorporates ‘read depth’ (i.e. the total number of reads covering that DNAm site), and ‘DNAm value’ (i.e. the proportion of methylated reads at that DNAm site). As with all sequencing applications, the total coverage, defined here as the total number of reads across the genome, is critical to the success of an experiment, as it will result in a higher average read depth at any individual DNAm point. Read depth influences both accuracy and statistical power. DNAm is measured as a proportion, therefore, when read depth is low there are only a finite number of possible values and the sensitivity of bisulfite sequencing is constrained. For example, a DNAm point covered by only four reads can only have five possible configurations of the ratio of methylated to unmethylated reads (4:0, 3:1, 2:2, 1:3, 0:4) resulting in the possible DNAm proportions of 0.00, 0.25, 0.50, 0.75, or 1.00. This lack of sensitivity has a direct effect on the magnitude and accuracy of differences that can be detected between groups, meaning that DNAm points with low average read depth may not have sufficient power for the detection of small or even moderate changes in DNAm. This is particularly pertinent as many studies of differential DNAm in complex phenotypes and disease typically identify changes of <5% [8, 18]; such small differences are likely to require precise proportions of the DNAm to be detected.

An additional challenge for the interpretation of bisulfite sequencing data compared to array-based methods, which have a fixed content, is that the precise regions of the genome covered by sequencing reads generated in any given experiment can be highly variable. This means that DNAm sites captured in a sequencing experiment may not contain many DNAm points, and that even where the DNAm points have been assayed across many of the samples, the read depth is potentially highly variable. This results in a matrix of DNAm values with a high proportion of missing data, effectively lowering the sample size at that DNAm site, in turn reducing the power to detect associations in analysis.

The gold standard bisulfite-sequencing method is whole genome bisulfite sequencing (WGBS) [19], although this can be cost prohibitive for many studies and is not yet amenable for large epidemiological analyses. Furthermore, in a study where the main interest is cytosines, in particular at CpG sites, a high number of WGBS reads are uninformative. Reduced representation bisulfite sequencing (RRBS), in contrast, involves a target enrichment step using the methylation-insensitive enzyme Mspl to target CpG-rich regions of the genome [20] prior to bisulfite conversion. This increases the proportion of informative sequencing reads, and RRBS typically interrogates DNAm sites in 85-90% of CpG islands [21, 22].

While multiple tools exist for the alignment and quantification of DNAm from bisulfite-sequencing data (e.g. *Bismark* [17], *GSNAP* [23], *BSMAP* [24], *BS-Seeker3* [25]), there is no consensus about the optimal approach for determining the appropriate minimum read depth or number of DNAm points required to ensure high-quality data for a well-powered statistical analysis. For example, existing studies have utilized a huge variety of read depth thresholds; a relatively arbitrary value between 5-20 reads per DNAm point is often used in filtering steps [26–29], most commonly with no justification provided for the use of that threshold. There is also no consensus as to what to do with DNAm sites that have very few DNAm points. Part of this inconsistency arises from a lack of guidelines or studies exploring how read depth and missingness influence statistical power.

The aim of this study was to determine the relationship between read depth and the accuracy of DNAm quantification, as well as the effect of missing DNAm points on statistical power for identifying group differences in DNAm with a particular focus on RRBS studies. Using properties derived from a large RRBS dataset generated by our group, we designed a simulation framework to explore how accuracy changes as a function of read depth, as well as comparing the DNAm level estimated from RRBS data with levels quantified using a novel Illumina array [30]. We then extended our simulation framework to investigate how statistical power to identify differences in DNAm level between groups varies as a function of read depth and sample size while also considering the effect of i) the level of DNAm at individual DNAm sites, ii) the expected difference in DNAm between groups, and iii) the balance of sample sizes between comparison groups. Our data-driven approach highlights the importance of filtering by minimum read depth and minimum number of DNAm points per DNAm site, and illustrates how the choice of threshold is influenced by the specific study design and the expected differences between groups being compared. Finally, we present an approach for estimating statistical power for a bisulfite sequencing study for a given read depth and minimum DNAm points filtering threshold which can be used to improve the detection of true positives and reproducibility of findings. Our tool, POWer dEtermined REad Depth filtering for Bisulfite Sequencing (POWEREDBiSeq), is available at https://github.com/ds420/POWEREDBiSeq as a resource to the community.

## Results

### Read depth in RRBS data follows a negative binomial distribution, while the level of DNAm is bimodally distributed

As part of an ongoing study of aging, we profiled DNAm in 125 frontal cortex samples dissected from mice aged 2-10 months old using a modified version of the original RRBS protocol [20] (see **Methods**). Prior to quality control filtering, a mean of 41,199,876 (SD = 6,753,486) single end reads were generated per sample (**Additional File 2**). The quality of the sequencing data was assessed using *FastQC* [31], before reads were aligned to the *mm10* reference (GRCm38) genome using *Bismark* [17]. Here, we define DNAm sites as vectors, such that each DNAm site has a DNAm point per sample, containing read depth and DNAm values. That is, DNAm site = {DNAm point_1_ = {m_1_, rd_1_},…, DNAm point_i_ = {m_i_, rd_i_},…, DNAm point_n_ = {m_n_, rd_n_}}, for i in 1 to n samples, where mi represents the proportion of DNAm at a DNAm pointi, and rd_i_ is the read depth, defined here as the total number of reads at the DNAm point. If rd_i_ is 0, there will be no DNAm point associated with sample i. Across all samples, there was a total of 64,199,621 distinct DNAm points covered (including CpG, CpH and CHH sites), with a total of 3,419,677 different DNAm sites assayed, and each sample containing a mean of 2,170,454 (SD = 124,281) DNAm points across all DNAm sites. We characterized the distribution of read depth for each sample across DNAm points, observing a unimodal discrete distribution, skewed to the left and characterized by a long tail (**Figure 1A**). This distribution is typical of count data and is expected in sequencing datasets where the vast majority of DNAm points are covered by relatively few reads and a minority of DNAm points are covered by a large number of reads. Across all DNAm points, 22.1% (60,117,549) had less than or equal to than 5 reads and 3.30% (8,941,868) had more than 100 reads. Next, we visualized the distribution of DNAm levels across all DNAm points, observing the expected bimodal distribution, with the majority of DNAm sites being either completely methylated (50% of DNAm sites > 0.95) or unmethylated (49% of DNAm sites < 0.05) [32] (**Figure 1B**).

**Figure 1:**
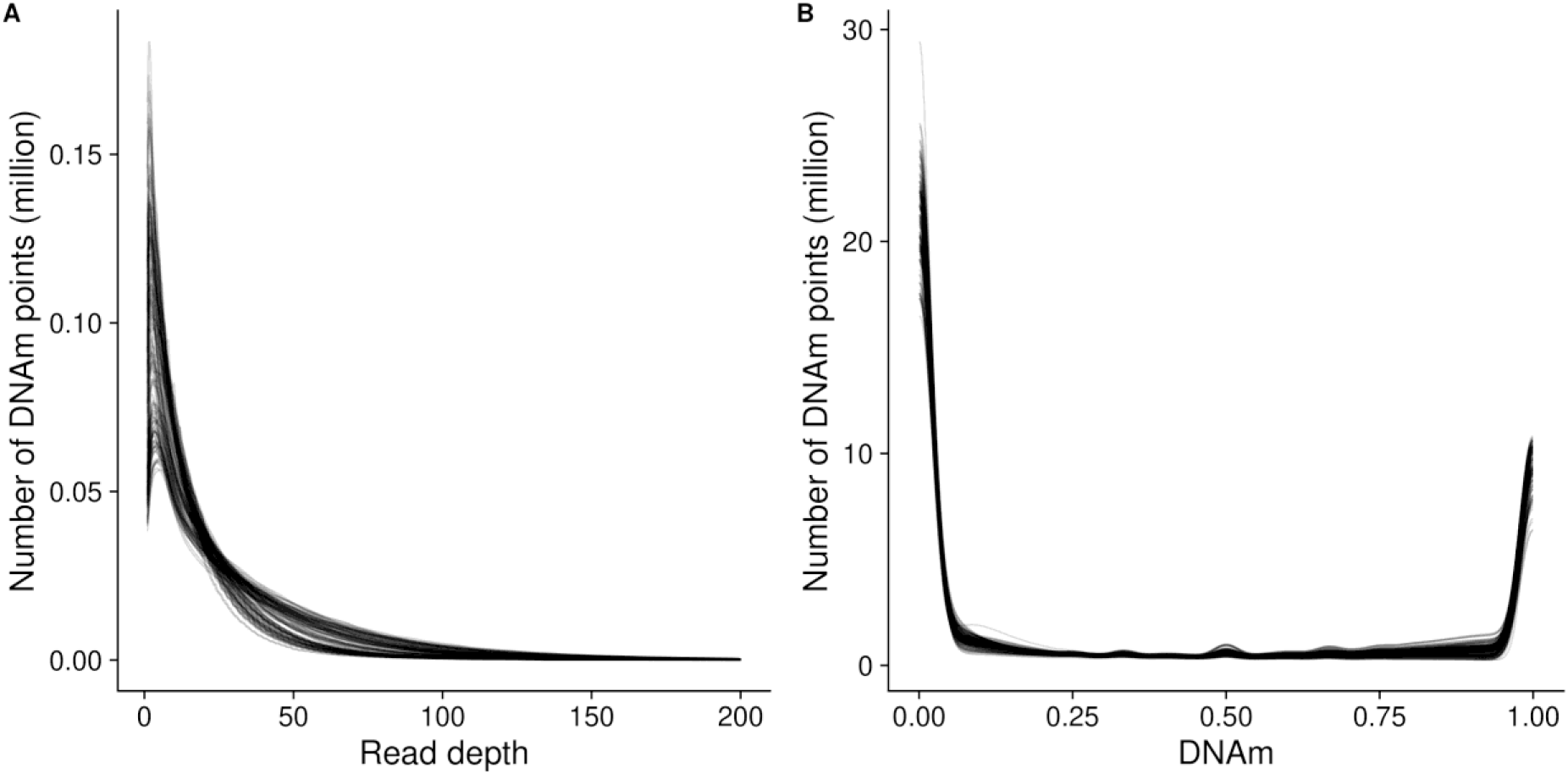
Characterization of read depth and mean DNAm across the DNAm points profiled by RRBS. The distribution of **A**) read depth across DNAm points and **B**) proportion of DNAm across DNAm points. Each line represents one sample. Read depth plots were capped at a read depth of 200 to facilitate the interpretation of plots, with less than 0.5% (1140174) of DNAm points being characterized by >200 reads.

### Read depth has a dramatic, non-linear effect on accuracy of DNAm estimates

One consequence of low read depth in RRBS data is reduced sensitivity for the quantification of DNAm at DNAm points. While DNAm points that are either completely methylated or unmethylated can theoretically be characterized precisely with a single read, this is not the case for DNAm points with intermediate levels of DNAm, which may be inaccurately classed as methylated or unmethylated at low read depths. To understand the extent of this problem, we compared the proportion of DNAm values at extremes (less than 0.05 or greater than 0.95), with increasing read depths across DNAm points (**Figure 2A**). As expected, the proportion of DNAm sites estimated to have extreme levels of DNAm was greater at lower read depths; 86.1% (SD = 4.94) of sites were estimated to have DNAm >0.95 or <0.05 at a read depth of 5, compared to 64.7% (SD = 6.90) at a read depth of 50. This suggests that, compared to DNAm points with a read depth of 50, more than 20% of DNAm points with a read depth of 5 may have been inaccurately classified as having an extreme level of DNAm.

**Figure 2:**
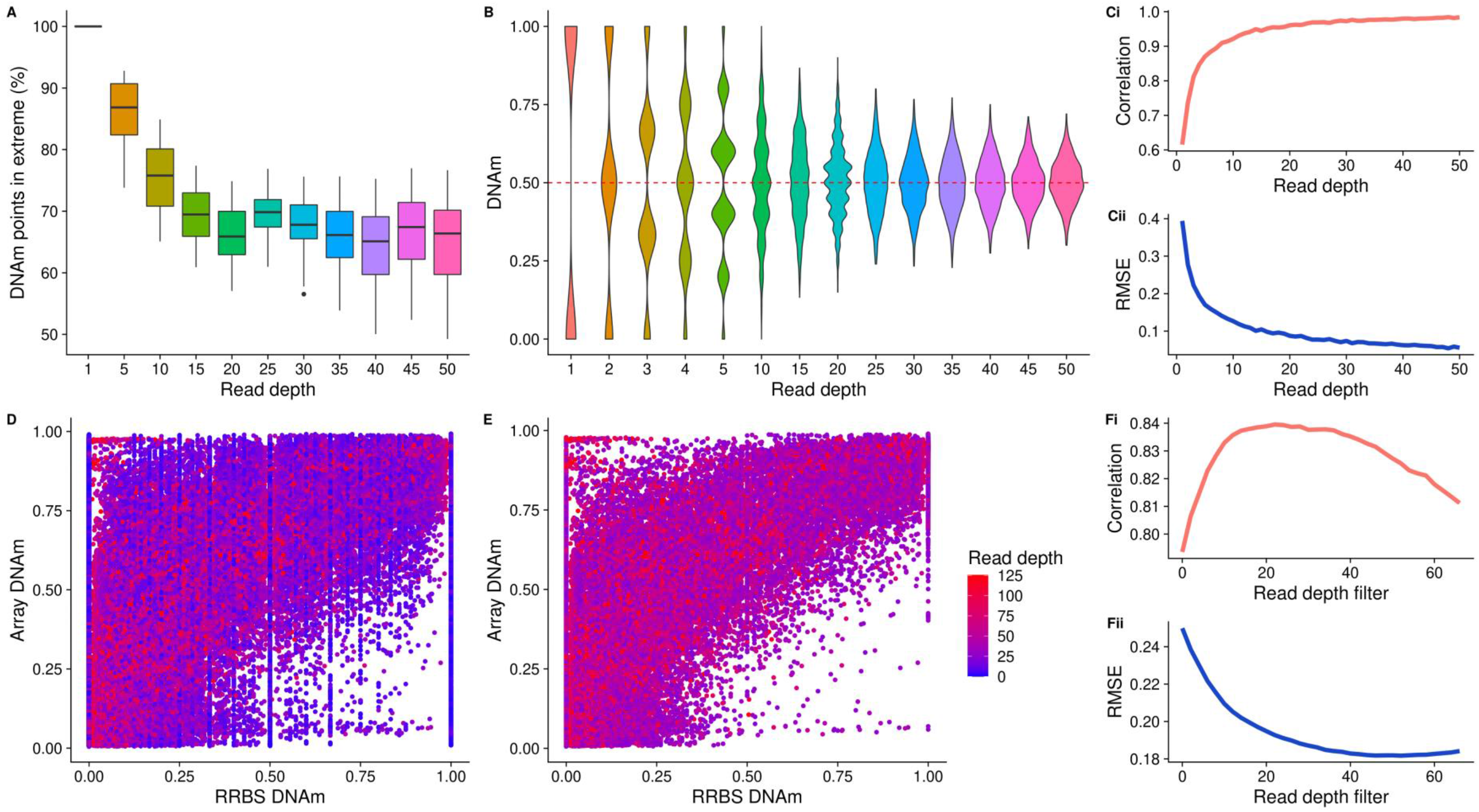
The consequence of ‘missingness’ in RRBS data demonstrated by array and simulation bisulfite-sequencing data. **A**) A boxplot showing the proportion of DNAm points that have ‘extreme’ DNAm (0.05 < DNAm < 0.95) calculated for DNAm points with different read depths (x axis). **B**) Violin plots showing the distribution of estimated DNAm values from a simulated bisulfite sequencing experiment for a DNAm site where the true value is 0.50, as a function of read depth. Line graphs showing the Pearson correlation (**Ci**) and root mean squared error (RMSE) (**Cii**) between simulated and ‘real’ DNAm values for 1000 DNAm points as a function of read depth. These analyses used a subset of real data selected to contain DNAm points with read depth >10 and evenly distributed DNAm (see **Methods**). Scatterplots of DNAm values quantified using RRBS (x-axis) and a custom vertebrate Illumina DNAm array [30] (y-axis) in matched samples (n = 80) for **D**) all DNAm points and **E**) the subset of DNAm points with read depth greater than the peak Pearson correlation read depth in **Fi** (i.e. 22 reads). Line graphs showing **Fi**) the Pearson correlation and **Fii**) error (RMSE) of RRBS data and array data as a function of the read depth filter applied to the RRBS dataset.

To formally quantify the error in estimating DNAm, we used simulations of increasing read depth to estimate DNAm for a hypothetical DNAm site with an intermediate level of DNAm (0.50), calculating the difference between the estimated and true DNAm level. For read depths <10, we observed a discrete distribution of estimated DNAm (Figure **2B**), with the range of predictions spanning 0.00 – 1.00 but centered on 0.50. In line with the Central Limit Theorem, we observe that as read depth increases, the distribution of estimated DNAm levels becomes more continuous and normally distributed around a DNAm value of 0.50. We expanded these simulations to consider DNAm sites with DNAm levels across the full distribution of possible values. We simulated 10,000 DNAm points with DNAm uniformly sampled between 0.00 – 1.00 and sampled 10,000 RRBS DNAm points with matched DNAm levels for comparison (see **Methods**). We found that as read depth increases, the correlation across DNAm points between estimated and actual DNAm level tends towards 1.00 (**Figure 2Ci**) and the RMSE tends towards 0.00 (**Figure 2Cii**). However, these effects are non-linear, with more dramatic improvements in accuracy occurring at lower read depths; i.e. there is a jump from a correlation of 0.589 to 0.926 between 1 and 10 reads with relatively minimal gains after that. Similarly, the RMSE drops from 0.404 at a read depth of 1.00 to 0.124 at a read depth of 10.

### RRBS and Illumina arrays DNAm values correlate highly

Commercial DNAm arrays, such as the Illumina EPIC BeadChip array, are commonly utilized as an alternative strategy to bisulfite sequencing approaches in large human studies, due to their relatively low cost and the ease of interpreting data [33]. To further characterize the accuracy and sensitivity of RRBS, we performed a comparison with DNAm levels quantified using a novel Illumina Beadchip vertebrate DNAm array [30] on an overlapping set of 80 mouse frontal cortex DNA samples. A total of 3,552 unique DNAm sites were quantified in both the RRBS and array datasets, with each RRBS sample containing a mean of 2,263 overlapping DNAm data points (SD = 104). First, we compared the distribution of DNAm estimates across all DNAm points between the two technologies, observing the expected bimodal distribution with both approaches (**Supplementary Figure 1**). Of note, the array data contains a higher proportion of DNAm sites with intermediate levels of DNAm (0.05-0.95), and the unmethylated and methylated peaks are shifted inwards from the boundaries, highlighting the reduced sensitivity of the array for quantifying extreme levels of DNAm [16]. In contrast, the peaks in the RRBS data are at 0.00 and 1.00. The array samples also have less variability between samples, with distributions looking nearly identical, due to DNAm points being consistently characterized for each DNAm site. Directly comparing the estimated level of DNAm between the two assays, we observed a strong positive correlation (Pearson correlation = 0.794) even with no read depth filtering in the RRBS data (**Figure 2D**). The correlation between assays increases as more stringent read depth filtering is applied to the RRBS data, with the maximum correlation (Pearson correlation = 0.840) obtained at a read depth threshold of 22 (**Figure 2E, Fi**). Although this correlation indicates a relatively strong relationship between the estimates of DNAm quantified using RRBS and the Illumina array, it does not necessarily indicate that the DNAm estimates generated by the two platforms are equal. Closer inspection showed that the relationship between RRBS- and array-derived DNAm estimates is not linear (**Figure 2D**), and therefore we also explored absolute differences in DNAm estimates between the two assays. We observed a notable skew, with DNAm estimates from the array being generally higher than those from RRBS (mean difference = 0.112, SD = 0.223), and this relationship was observed regardless of read depth (**Supplementary Figure 2**). As expected, the root mean squared error (RMSE) between DNAm estimates generated using array and RRBS decreases as the stringency of read depth filtering in the RRBS dataset increases (**Figure 2Fii**), plateauing at a read depth of ~30. Of note, the minimum RMSE observed was 0.180, suggesting some systemic differences between the two platforms in estimated DNAm levels.

### A subset of DNAm sites have consistent read depth across DNAm points

In order to perform a statistical analysis of DNAm differences between groups (e.g. in a study of cases vs controls), multiple samples, usually representing biological replicates, are required. We have demonstrated the importance of filtering RRBS data by read depth on obtaining accurate estimates of DNAm, however, this has the consequence of increasing the number of missing DNAm points (**Figure 3A**). Of note, we found that read depth is not random across DNAm sites, but highly correlated between pairs of samples (**Figure 3B**). To demonstrate this further, we iteratively increased the number of samples and calculated the proportion of DNAm points shared across DNAm sites (**Figure 3C**). The proportion of DNAm points present decreases in a non-linear manner before plateauing at 0.20, demonstrating that there is a subset of DNAm sites for which read depth is greater than 0 across all or most DNAm points. DNAm sites containing all possible DNAm points, that is, each DNAm point had a read depth >1, were found to have consistently higher read depth, with a strong correlation in read depths between DNAm points (**Figure 3D**). We hypothesized that the correlation in read depth between samples resulted from the enrichment strategy used in RRBS, meaning that specific CpG-rich regions are dramatically overrepresented in the sequencing data across all samples, and as expected, the common DNAm sites containing all possible DNAm points were enriched in CpG islands compared to all DNAm sites (**Figure 3E**).

**Figure 3:**
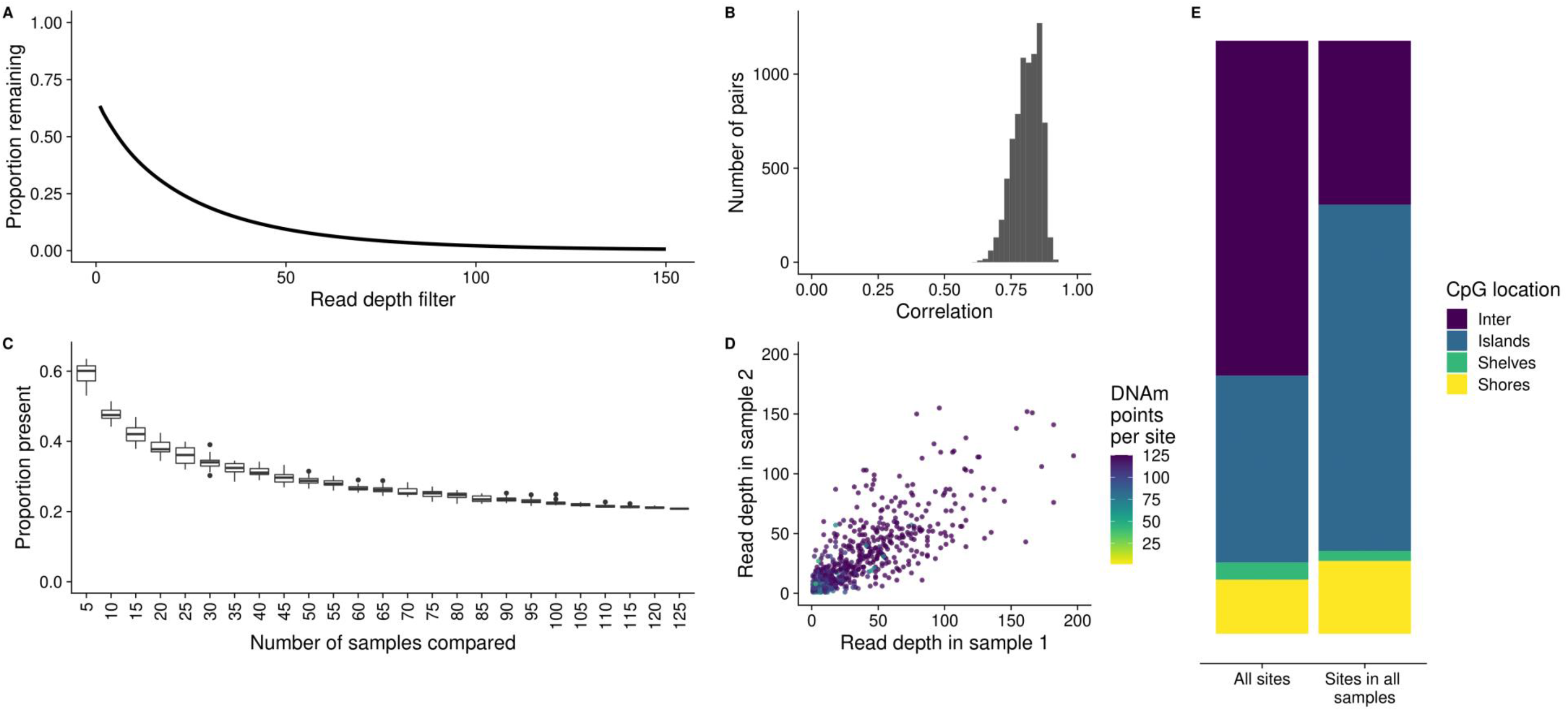
A subset of higher read depth DNAm sites are over-represented in RRBS datasets. **A**) A line graph of the mean proportion of DNAm sites remaining (y-axis) after filtering by increasing read depth thresholds (x-axis). **B**) The Spearman’s correlation of read depth between all pairs of samples. **C**) The proportion of overlap in the DNAm points present across an increasing number of samples compared. **D**) Read depth plotted from two randomly selected samples, colored by the number of DNAm points that the DNAm site that have a read depth > 0. 1000 DNAm points were randomly selected and read depth is plotted up to 200 to facilitate the interpretation of plots. **E**) The proportion of DNAm sites in intergenic regions (purple), CpG islands (blue), shelves (green) and shores (yellow) for all DNAm sites and all DNAm sites with read depth >1 across all samples.

### Simulated data demonstrates the consequence of read depth, sample size, and mean DNAm difference per group on power

Statistical power to identify differences in DNAm between two groups (e.g. cases vs controls), defined as the proportion of successfully detected true positives, will vary across DNAm sites and is influenced by multiple variables. In bisulfite sequencing studies, these include read depth, the number of samples in each group, the ratio of group sizes, the mean DNAm level, and the expected difference in DNAm between groups. We explored how each of these variables influences power by simulating bisulfite sequencing data for a given DNAm site following the framework laid out in **Supplementary Figure 3**. Briefly, a two group comparison was simulated, with sample size, mean read depth, μDNAm (the mean DNAm across the DNAm point) and ΔμDNAm (the mean difference in DNAm between groups) used as input variables that were either kept constant or varied to observe the effect on power. Each exemplar DNAm site was simulated 10,000 times, containing all DNAm points for the given sample size. A two-sided t-test was used to compare groups and power calculated as the proportion of p-values smaller than 5×10^−6^. It is important to note that all parameters, including r, the p value threshold for power, and number of DNAm sites simulated, were selected with the aim of visualising how power might change with each variable in turn. Subsequent findings are based on exemplar DNAm sites, and exact values should be taken as such; they may not be representative of a wider study, as our aim was solely to characterize the relationship between each variable and statistical power. The values used to generate the results for each variable shown in **Figure 4** can be found in **Supplementary Table 1**.

**Figure 4:**
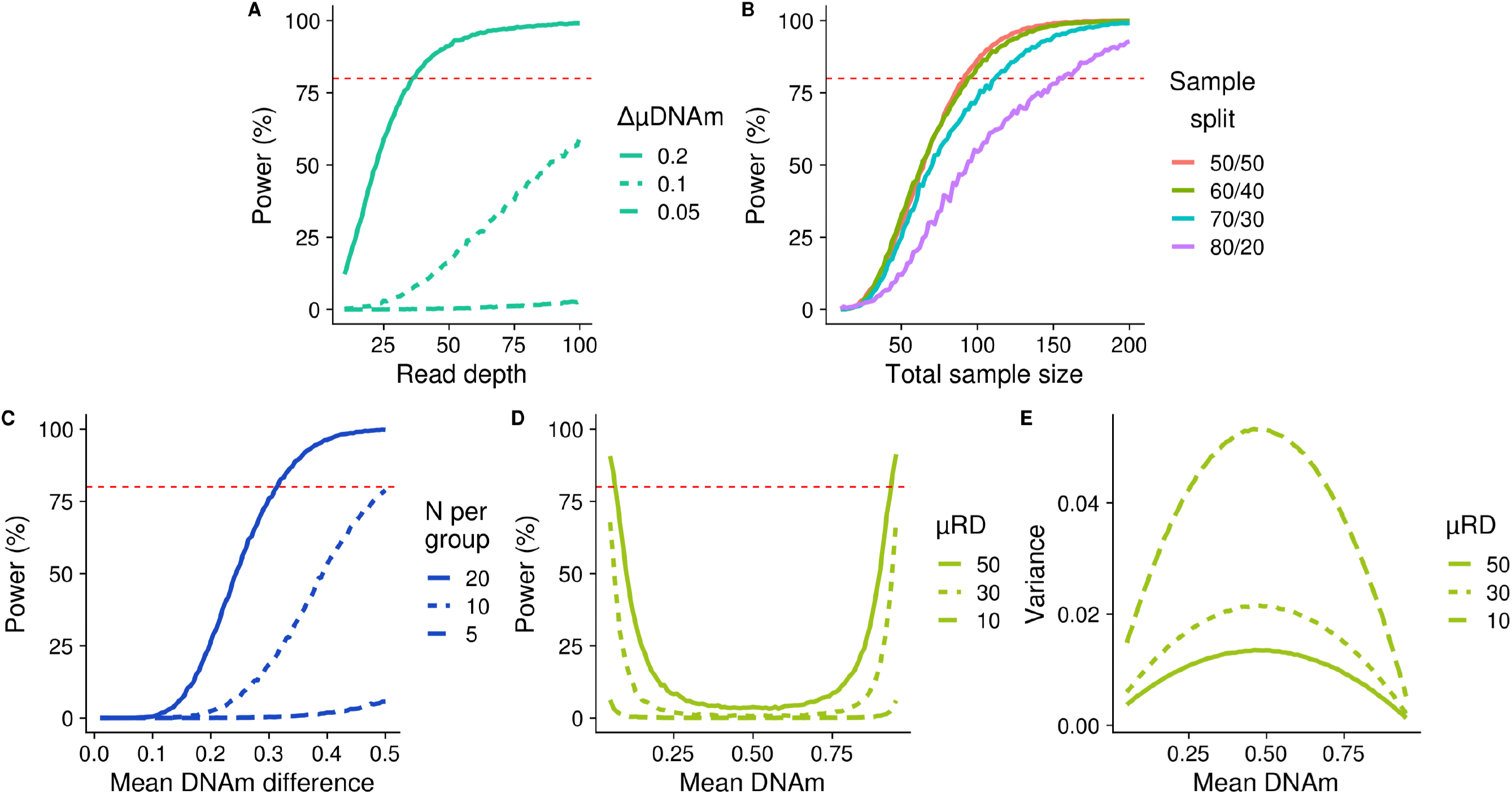
Power is influenced by read depth, sample size, and mean DNAm level in two-group comparisons. Power curves plotting statistical power to detect significant differences in DNAm between two groups as a function of **A**) read depth, **B**) sample size and the effect of an unbalanced sample size between groups, **C**) the mean difference in DNAm between the groups and **D**) the mean DNAm at simulated DNAm sites. **E**) The variance for the simulated data shown in panel **D**. Simulations were performed 10,000 times with a negative binomial parameter of r = 1.5.

As expected, increased read depth had a positive effect on power across each of the scenarios we considered, however, the potential gains are highly dependent upon the specific combination of parameters (**Figure 4A**). For example, in a scenario where each group contains 30 samples and the mean DNAm level is 0.25, there is a relatively dramatic increase in power to detect a DNAm difference of 0.20 between groups as read depth increases, with 80% power at a mean read depth of 37, although there are minimal gains with read depths > 50. In contrast the gain in power with increased read depth is much less pronounced when detecting a mean DNAm difference of 0.10, and there is very little power at any read depth to detect a DNAm difference of 0.05. Therefore, if small effect sizes are relevant for the phenotype under study, power will need to be increased through other methods, e.g. increased sample size, as read depth filtering alone will not be sufficient.

We next investigated the effect of sample size and the ratio of group sizes on power (**Figure 4B**), concluding that the optimal design in terms of maximizing power is to have equal sized comparison groups, assuming that the total sample size is constant. Fixing mean read depth to be 20 and a mean DNAm level of 0.25, our simulations showed that to have 80% power to detect a DNAm difference of 0.20 between groups a total sample size of 94 is required when the sample size ratio between groups is 60:40 (56 and 38 samples, respectively), which is only two more samples than required when the sample size ratio is balanced (i.e. 50:50). In the most extreme scenario we considered, an 80:20 ratio between groups, a total of 154 samples (123 and 31, respectively) are needed to have 80% power to detect a DNAm difference of 0.20 between groups. This has implications for the handling of DNAm sites where DNAm points are missing; it suggests that there may be a tolerable level of ‘missingness’ when comparing DNAm between groups that can be ‘rescued’ by having a greater sample size in the second comparison group. As with read depth (**Figure 4A**), we found a non-linear relationship with power for both sample size (**Figure 4B**) and mean DNAm difference between groups (**Figure 4C**). Where each of these variables is the limiting factor, we found that the greatest gains in power occurred initially, with diminishing returns at higher levels and an eventual plateau. Where other variables act to reduce the overall power, the power curve is flattened and a plateau is not reached. One interesting observation from our simulations was the U-shaped relationship between power and mean level of DNAm at a given site (**Figure 4D**). Power is highest at DNAm sites with either very low or very high levels of DNAm, and decreases to a minimum at intermediate levels of DNAm. We hypothesize that this reflects the relationship between the mean and variance in DNAm [34] (**Figure 4E**), where the variance is lowest at the extremes, an artefact of DNAm being measured as proportion bounded at 0.00 and 1.00.

### Simulated bisulfite sequencing studies can be utilized to estimate power given suggested filtering

Our results indicate that, given the complex interplay of multiple experimental parameters, the choice of threshold for filtering DNAm sites is not always straightforward and will depend on the specific research question being addressed. Furthermore, the power calculations presented so far only consider a single DNAm site, whereas genome-wide comparisons of DNAm typically involve the analysis of hundreds of thousands of DNAm sites; given the effect of the properties of DNAm sites (e.g. in mean DNAm level) on power, no DNAm site can be considered to be ‘representative’ of the others. Therefore, we extended our simulation framework to quantify a study-level power statistic that considered all DNAm sites, allowing for the calculation of power given an RRBS dataset, and the read depth and minimum DNAm points per DNAm site filtering to be carried out. The extension of the simulation framework can be seen in **Supplementary Figure 4** and is described in **Methods**. Briefly, an actual RRBS data set was used to estimate the simulation parameters (namely, sample size, μDNAm, μRD and negative binomial parameter, r) so that simulations reflect the real data. We compared the real and simulated data finding that the distribution of simulated read depths is highly comparable to real data for lower read depths (**Figure 5Ai**). Higher read depths do not seem to be captured as accurately by the negative binomial distribution, however, given that 95% of DNAm points have read depth < 85 (**Figure 5Aii**), this should be less important to the simulation. Overall, simulated DNAm estimates were similar to real DNAm levels across DNAm points, although there was some deviation, for example, a slight under representation of DNAm points with DNAm proportions above 0.25 and an overrepresentation of DNAm points with DNAm proportions above 0.25 (**Figure 5B**).

**Figure 5:**
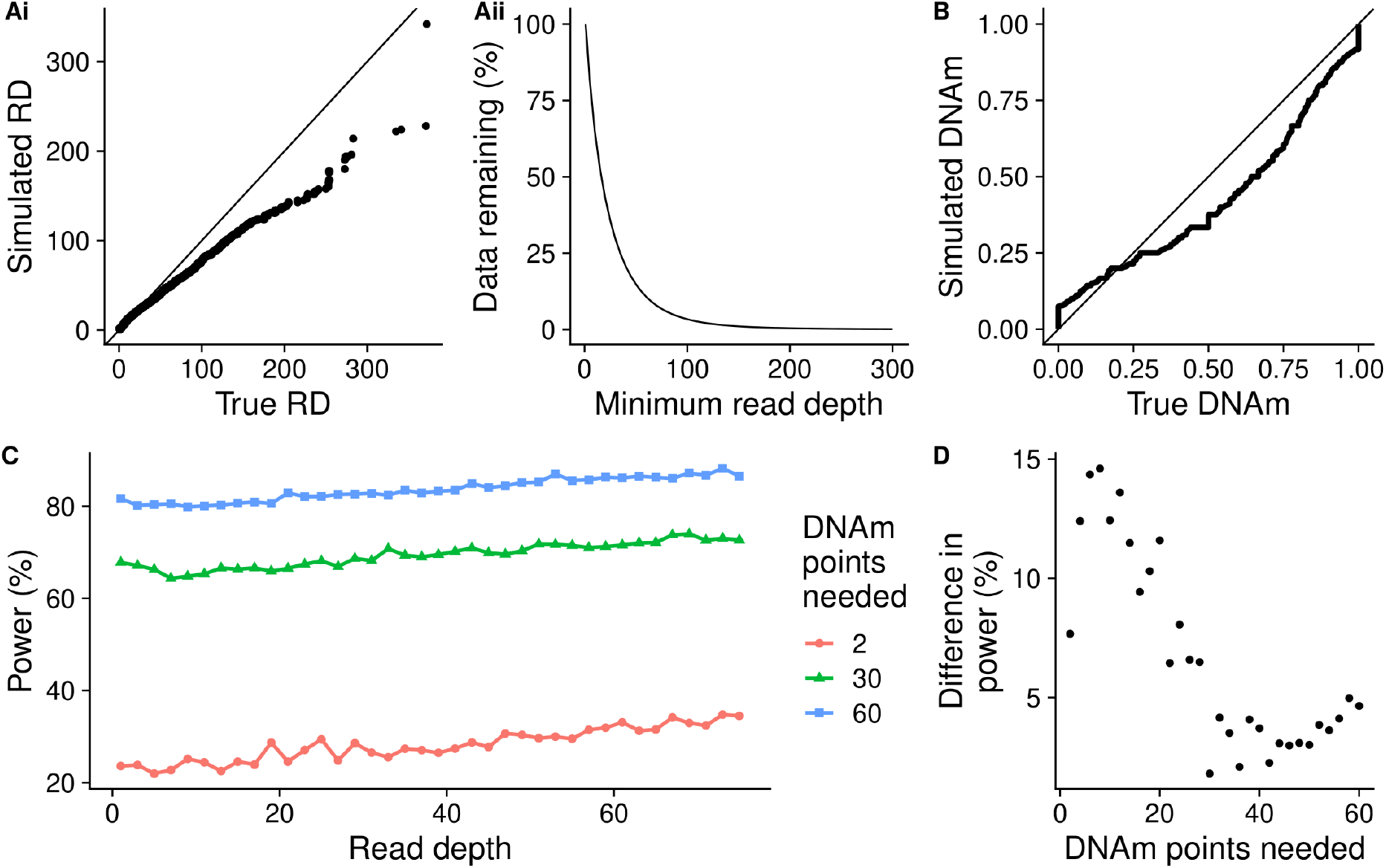
Summarizing the simulation and predictions of POWEREDBiSeq. **Ai**) A QQplot comparing the read depth of 10,000 simulated DNAm points to 10,000 randomly sampled true DNAm points from an RRBS dataset. **Aii**) The proportion of DNAm points remaining in the RRBS dataset with read depths >x. **B**) A QQplot comparing the DNAm of 10,000 simulated DNAm points to a 10,000 randomly sampled true DNAm points. **C**) The relationship between the difference in power predicted by POWEREDBiSeq at different minimum sample sizes (n = 2, 30 and 60) as the minimum read depth threshold is increased, with a mean difference between groups of 0.06. **D**) The relationship between the increase in power to detect a mean difference in DNAm between groups of 0.06 predicted by POWEREDBiSeq at a read depth of 75 compared to power at a read depth of 1 as a function of the number minimum of samples per group.

To demonstrate the methodology, we considered a hypothetical study design with a total of 125 samples, specifying an expected mean DNAm difference between groups of 0.06, picked arbitrarily to allow for power visualization. To profile how read depth influences the power of the study, we incrementally increased the minimum read depth from 1 to 75, and to investigate the effect of the minimum number of DNAm points needed to find a difference between groups, we chose three arbitrary values: 2, 30 and 60. Power only increased subtly as read depth filtering became more stringent (**Figure 5C**), compared to the gain of increasing the number of DNAm points. However, the gain is not consistent across all study designs, with greater gains in smaller studies (**Figure 5D**). For example, with a minimum of two DNAm points per group, increasing the read depth threshold from 1 to 75 resulted in an increase in power of 10.9%, compared to a smaller increase of 4.83% and 4.89%, respectively, when the minimum DNAm points were set at 30 or 60. Our analysis reaffirms the interplay between all study-specific experimental variables. However, it should be noted that even with the most extreme read depth filtering, the maximum power for a group with a minimum of two DNAm points is still dramatically lower that the power of a study with a larger minimum and no or negligible filtering. Finally, we summarized our study wide power calculation in the R function POWer dEtermined REad Depth filtering for Bisulfite Sequencing (POWEREDBiSeq), which is available as a resource to the community at https://github.com/ds420/POWEREDBiSeq. The calculation results in largely consistent and normally distributed predictions of power, however, outliers can occur, suggesting that multiple iterations should be performed (**Supplementary Figure 5**).

## Discussion

In this paper, we systematically characterize the properties of a representative RRBS dataset, assessing the distribution of read depth and missing data across DNAm sites. Using our framework of bisulfite sequencing data simulation, we investigate the impact of various study variables (e.g. read depth, group size, skewness in group size, and magnitude of DNAm difference) on the accuracy of DNAm quantification, and power to detect DNAm differences between two groups. As a resource to the community, we have developed a tool (POWEREDBiSeq), which utilizes our findings to predict power for individual study designs, accounting for the filtering to be applied.

When comparing to simulated data, we found that the accuracy to detect a given DNAm difference between groups improves with increased read depth. This likely reflects the fact that count data is only able to represent continuous data if the number of counts (i.e. sequencing reads) is high enough. Overall, we found a strong correlation in DNAm estimates derived from RRBS and Illumina DNAm array data; this relationship increases with minimum read depth filtering and reaches a maximum when excluding DNAm sites covered by less than 22 reads. The high correlation between platforms and the relationship with read-depth concurs with previous analyses comparing RRBS and Illumina array in human samples [35]. This finding has implications for studies using RRBS to identify differences in DNAm as it highlights the importance of read depth filtering in generating an accurate measure of the true DNAm level.

We investigated the impact of various experimental variables on power, defined as the proportion of true positives detected in a two-group comparison, in a bisulfite-sequencing study utilizing simulated data. We observed that these variables (read depth, sample size, DNAm difference between groups and mean DNAm at a given DNAm site) act together to influence power. Read depth, sample size and DNAm difference between groups will all limit power in a certain range, with power plateauing at 100% when they are no longer the limiting factor. DNAm level at a DNAm site has a U-shaped relationship with power, where DNAm points with extreme DNAm (near 0 and 1) are more powered to identify between-group differences primarily because the variance in DNAm at these DNAm sites is smaller. Our findings highlight the importance of data filtering for maximizing power; the minimum number of DNAm points needed across each DNAm site to be compared has a dramatic effect on power, as it dictates the minimum effective sample size at any one DNAm site. Read depth also influences power, although we observed that read depth filtering alone cannot overcome an inadequate study design (i.e. too few samples). As a resource to the community, we have summarized our data simulations so that others can apply them to their data to calculate the power to identify between-group differences in DNAm within the context of their specific study design. Our scripts are packaged into the POWEREDBiSeq application (https://github.com/ds420/POWEREDBiSeq) which allows users to optimize their power by, for example, simulating the effects of increasing their sequencing read depth filtering threshold or minimum DNAm points across groups.

Although our analyses and simulations focused on RRBS datasets, many of our conclusions are valid for other types of bisulfite sequencing data. For example, the relationship between read depth and accuracy applies to any bisulfite sequencing based DNAm experiment that profiles DNAm at a single nucleotide resolution. Additionally, the relationship between power and read depth, sample size, DNAm difference, and mean DNAm is also relevant for other sequencing based DNAm studies. Various methods differ in read depth and the distribution of DNAm sites sequenced across the genome. Targeted bisulfite sequencing (TBS), for example, typically profiles a more restricted set of DNAm sites than RRBS, as only regions of interest are enriched. This results in a more uniform distribution of reads across DNAm points, which acts to improve power across the study. In whole genome bisulfite sequencing (WGBS) studies, however, while more DNAm sites are interrogated across the genome as a whole, the read depth per DNAm point tends to be lower than that obtained using RRBS or TBS. POWEREDBiSeq can be applied to other bisulfite sequencing types because the internal variables, such as DNAm distribution and number of DNAm sites, are calculated based on input data. For the same reason, POWEREDBiSeq is also applicable to DNAm at CHH and CGH sites, which are often covered in bisulfite sequencing studies but have dramatically different properties to DNAm at CpG sites, although it is important to verify that the simulated and real data distributions are alike. In datasets with a frequent occurrence of high read depths across DNAm points (>100), some caution in the use of POWEREDBiSeq is warranted, as we found that the negative binomial distribution underestimates higher read depths when simulating data. This was not pertinent in our case as the 95% of sites had a read depth below 85.

The results of POWEREDBiSeq will be dependent on the planned filtering stringency of the user, as well as the biological question that the bisulfite sequencing experiment aims to address; for example, a study looking into DNAm changes between cancer and non-cancer samples will have higher power due to the comparatively large DNAm differences between groups [36] compared to those observed in many complex disease case and control studies [8, 18]. Bisulfite sequencing data generated in cell lines and genetically identical mouse models will be comparatively less ‘noisy’ than analyses of diverse human populations using heterogeneous tissues such as blood, resulting in increased power. Retaining poor quality (i.e. low read depth) DNAm sites in a bisulfite sequencing dataset increases the multiple testing burden, meaning it will be harder to identify true between-group differences in DNAm at higher quality, more adequately powered, DNAm sites. A limitation of POWEREDBISEQ and our data simulations is that they are based on a two-group comparison (e.g. cases vs controls), meaning our findings are not specifically applicable to more complex study designs. One question not addressed by our analysis is whether, for a given amount of available resource, it is optimal to sequence more samples at the same level or increase sequencing depth for a smaller number of samples. To explore this further, data from additional RRBS studies with a sufficient range of coverage would be needed.

To our knowledge, this is the first attempt to develop recommendations for bisulfite sequencing experiments based on sequencing read depth, minimum number of DNAm points and statistical power. We believe findings from this work will improve the reproducibility of bisulfite sequencing studies; we encourage researchers working in this field to clearly detail any data filtering steps and ensure an appropriate filter for read depth and other parameters has been applied, with justification for the choice of threshold.

## Methods

### DNAm quantification by RRBS

Genomic DNA was isolated from mouse cortex [37] using the AllPrep DNA/RNA Mini Kit (QIAGEN) and assessed for quality and quantity using the NanoDrop 8000 spectrophotometer (Thermo Fisher Scientific) and the Qubit high sensitivity assay (Qubit dsDNA HS Assay, Thermo Fisher Scientific). RRBS libraries were prepared using the Premium RRBS kit (Diagenode). The final RRBS library pools were distributed across thirty-two HiSeq2500 (Illumina) lanes and subjected to 50 bp single-end sequencing as previously described [20].

### Preprocessing the dataset

RRBS sequencing quality was assessed using *FastQC* (version v0.11.7) [31] with all samples characterized by high quality base calls (quality score >28 across all bases). Sequences were trimmed using *TrimGalore* (version 0.4.4_dev) [38], with a quality score of 20 and an error rate of 0.2 used to remove poor quality bases at the ends of reads. Reads with fewer than 20 base pairs after trimming were then removed. Reads were aligned to the mm10 (GRCm38) mouse genome [39] using *Bismark* v0.19.0 with default parameters [17], which implements *SAMtools* 1.8 [40] and *Bowtie2* v2.3.4.1 [41]. The total number of aligned reads and cytosines can be found in **Additional File 2**.

### Statistical methods

All subsequent analysis was carried out in R 3.5.2 (2018-12-20)[42] using the R packages *ggplot2* (version 3.2.1)[43], *Cowplot* (version 1.0.0) [44], *Tidyr* (version 1.0.0) [45], *Viridis* (version 0.5.1), *viridisLite* (version 0.3.0) [46], *colortools* (version 0.1.5) [47], and *reshape2* (version 1.4.3)[48].

### Annotating RRBS to the CpG islands

R packages *annotatr* (version 1.8.0)[49] and *GenomicRanges* (version 1.34.0)[50] were used to annotate CpGs to features for the analyses shown in **Figure 3E**. The *annotatr* package assigned CpG islands as per the mm10 reference annotation, with CpG shores defined as 2Kb upstream/downstream from the ends of the CpG islands, and CpG shelves as another 2Kb upstream/downstream of the farthest upstream/downstream limits of the CpG shores. The remaining genomic regions make up the inter-CGI annotation.

### DNAm quantification quantified by array

A subset of 80 DNA samples were additionally profiled using a custom Illumina DNAm array (the “HorvathMammalMethylChip40” [30]). Briefly, this array includes ~36k CpGs that are located in genomic regions highly-conserved across 50 mammalian species. Data was loaded from idat files into an RGChannelSet object using the *minfi* package (version 1.28.4) [51–57] and processed through the following steps: 1) checking the methylated and unmethylated intensities and excluding samples < 800, 2) confirming successful bisulfite conversion excluding samples with low conversion rates (<80%), 3) confirming correct sex using profiles from the X chromosome, and 4) confirming tissue type, excluding any sample predicted incorrectly based on DNAm profile. Prior to analysis data was normalised using the *Sesame* package (version 1.4.0)[58], and filtered to DNAm sites classed as mapping uniquely to the mouse genome, leaving 23,633 DNAm sites.

### Framework for simulating RRBS data

We developed an analytical framework to profile the power of RRBS DNAm sites, enabling us to vary different parameters such that we could explore a number of research questions. The DNAm site-level simulation workflow is described in **Supplementary Figure 3**, which aims to compare the DNAm between two groups, A and B. For each DNAm site simulated, there are 8 parameters to consider: *N_1_* and *N_2_* are the sample size each group, respectively, *μRD* is the mean read depth of the DNAm site to be simulated, *r* is a negative binomial parameter, described in more detail below, *μDNAm* is the mean DNAm across the DNAm site, *ΔμDNAm* is the mean difference in DNAm between groups, *nSites* is the number of DNAm sites to be simulated, and *pValue* is the p-value used to assess power.

When simulating a DNAm site, the first step is to simulate read depth. Read depth could be assigned an arbitrary value, or, where realistic variation across DNAm points was required, could be sampled from a negative binomial distribution [59]. The negative binomial distribution is defined by the parameters *r* and *p*, although within the R function *rnbinom()* can be defined by *μRD* and *r*, which can calculated from real data using equation (6), the derivation of which is as follows:

The negative binomial equations are:

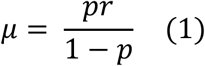

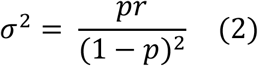

Where *μ* is the mean (in this case, *μRD*) and *σ*^2^ is the variance of the read depth data calculated across all samples. We want *r* in terms of *μ* and *σ*^2^. Multiply (2) by 1 − *p* and equate that and (1) to get:

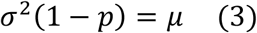

Rearrange for p:

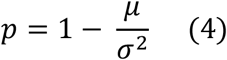

Substitute (4) into (1) and simplify:

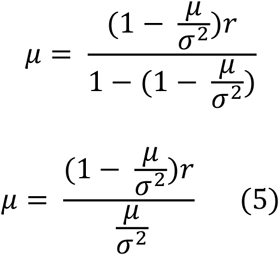

Rearrange (5) for *r*:

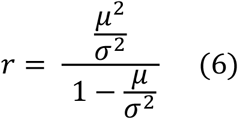

Once read depth was established, a binary value representing DNAm status was assigned to each read using the binomial distribution. For each read in group A, the probability of being methylated was *μDNAm*, and for group B was *μDNAm* ± *ΔμDNAm*, where the probability was bound between 0 and 1. The proportion of DNAm was calculated as the mean DNAm at each DNAm point.

The process was repeated for *nSites*. To calculate power, a two-sided t-test was performed between groups A and B. Power was defined as the proportion of DNAm sites for which the t-test p-value was smaller than *pValue*.

### Profiling the accuracy of RRBS data

To investigate how the distribution and accuracy of DNAm changed with increasing read depth, we considered a range of read depths (1 - 50). To profile accuracy across levels of DNAm in an RRBS study, we simulated 10,000 DNAm points per read depth, with DNAm sampled uniformly between 0 – 1. 10,000 DNAm points with matching DNAm were sampled from the RRBS data and correlation and RMSE were calculated between the true and the estimated DNAm points for each read depth.

### Profiling the power of RRBS data

To calculate the power of RRBS DNAm sites, we investigated a hypothetical two-group comparison study design (e.g. a case vs control analysis). We aimed to explore the effects of read depth, mean DNAm level, the sample size and sample size balance of groups, and the mean DNAm difference between groups on power. To this end, we utilized the simulation framework described above and in **Supplementary Figure 3** to simulate specific DNAm sites so that the resulting shift in power, given a change in a variable or combination of variables, could be visualized. The parameters assigned can be seen in **Supplementary Table 1**, where the variable parameter took a range of discrete values as seen in the x axes in **Figure 4**. *μRD* set was used as a negative binomial parameter, from which read depth (>0) was sampled. For group A, *μDNAm* was used as the probability of DNAm, sampled from the binomial distribution. For each set of parameters chosen, 10,000 DNAm sites were simulated. The *r* value was 1.5, and *pValue* 5×10^−6^, which were chosen arbitrarily to allow for the visualization of changing power.

### Profiling the power of RRBS studies given data filtering

We aimed to create a power calculator to determine the statistical power of a bisulfite sequencing study with specified read depth and minimum DNAm point filtering thresholds and specified mean DNAm difference between groups across a two-group study design. To this end, we utilized the simulation framework described above and in **Supplementary Figure 3** to simulate filtered data. The following input data was required (also described in **Supplementary Figure 4**): *RRBSTrue* - the unfiltered matrix of RRBS data, *ΔμDNAm* - the mean difference in DNAm between groups expected given the biology of the samples, *nDNAmPoint* - the minimum number of DNAm points needed per DNAm site, *RDFilter* – the minimum read depth filter to be applied, *pheno* – an optional variable dictating group membership.

These were used to estimate the variables for the framework in **Supplementary Figure 3**: *N_1_* and *N_2_* were assigned using *pheno*, or if *pheno* was not given, assigned as half of the number of samples in *RRBSTrue*. The data being simulated represented data that remained was post-filtering, therefore, given that we need at least *nDNAmPoint* DNAm points with sufficient read depth, *μRD* was calculated separately for the first *nDNAmPoint* DNAm points to the latter. For the first *nDNAmPoint* DNAm points, *μRD* was the larger of the mean read depth across *RRBSTrue* (estimated using 60,000 DNAm sites) and *RDFilter*, and subsequent read depth must be > *RDFilter*. For the remaining DNAm points, the mean read depth was used, where all simulated read depths < *RDFilter* were assigned a read depth of 0 to represent that they would get filtered out of the data. *r* was estimated using equation 6 and a subset of 60,000 DNAm sites from *RRBSTrue*. To estimate *μDNAm*, we first estimated the probability that a filtered DNAm site falls into one of the following ranges: 0-0.05, 0.05-0.95, 0.95-1, using a subset of 100,000 DNAm sites from *RRBSTrue*. The ranges were sampled using the probabilities calculated and a uniform distribution used to set *μDNAm* from the values across the selected range. To ensure that the subsets of *RRBSTrue* used to estimate variables were enough, we investigated the decline in prediction variability for each (**Supplementary Figure 6-8**).

40,000 DNAm sites were simulated, using the above inputs and step 1 of the workflow presented in **Supplementary Figure 3** and above. The resulting p-values were bootstrapped to result in the same number as the number of DNAm sites remaining in *RRBSTrue* after filtering by *RDFilter* and *nDNAmPoint*. The power was calculated using a Bonferroni correction for the number of DNAm sites remaining.

We created POWEREDBiSeq so that others can calculate their statistical power in bisulfite sequencing studies. The R function is available at https://github.com/ds420/POWEREDBiSeq.

## Declarations

## Ethics approval and consent to participate

All animal procedures were carried out at Eli Lilly and Company, in accordance with the UK Animals (Scientific Procedures) Act 1986 and with approval of the local Animal Welfare and Ethical Review Board.

## Consent for publication

NA

## Availability of data and materials

Data has been deposited in GEO under accession number GSE125957.

All scripts for this paper are publicly available and can be found in the Characterizing-the-properties-of-bisulfite-sequencing-data repository at https://github.com/ds420/Characterizing-the-properties-of-bisulfite-sequencing-data.

## Authors’ contributions

DSV, JM and EH wrote the manuscript, it was read and approved by all authors. IC performed the lab work. Analysis was led by DSV, who was assisted by AD.

## Acknowledgements

NA

## Additional File 1: Supplementary Figures

.docx file containing Supplementary Figures 1-10 and Supplementary Table 1

## Additional File 2: RRBS sample alignment and read depths

.xlsx file containing RRBS information on total number of reads aligned, unaligned ambiguously aligned, and total number of reads, as well as the number of methylated and unmethylated CpGs, CpH, and CHH’s, and total number of cytosines.

