## Additional File 1 for "Characterizing the properties of bisulfite sequencing data: maximizing power and sensitivity to identify between-group differences in DNA methylation"

**Supplementary Figures**

**Supplementary Figure 1: The distribution of DNAm levels across the genome profiled using RRBS or a custom array.** Density plot of the distribution of DNAm level across all DNAm sites, where each line represents one of the 80 overlapping samples profiled using a custom Illumina array [1] (blue)or RRBS (green). .

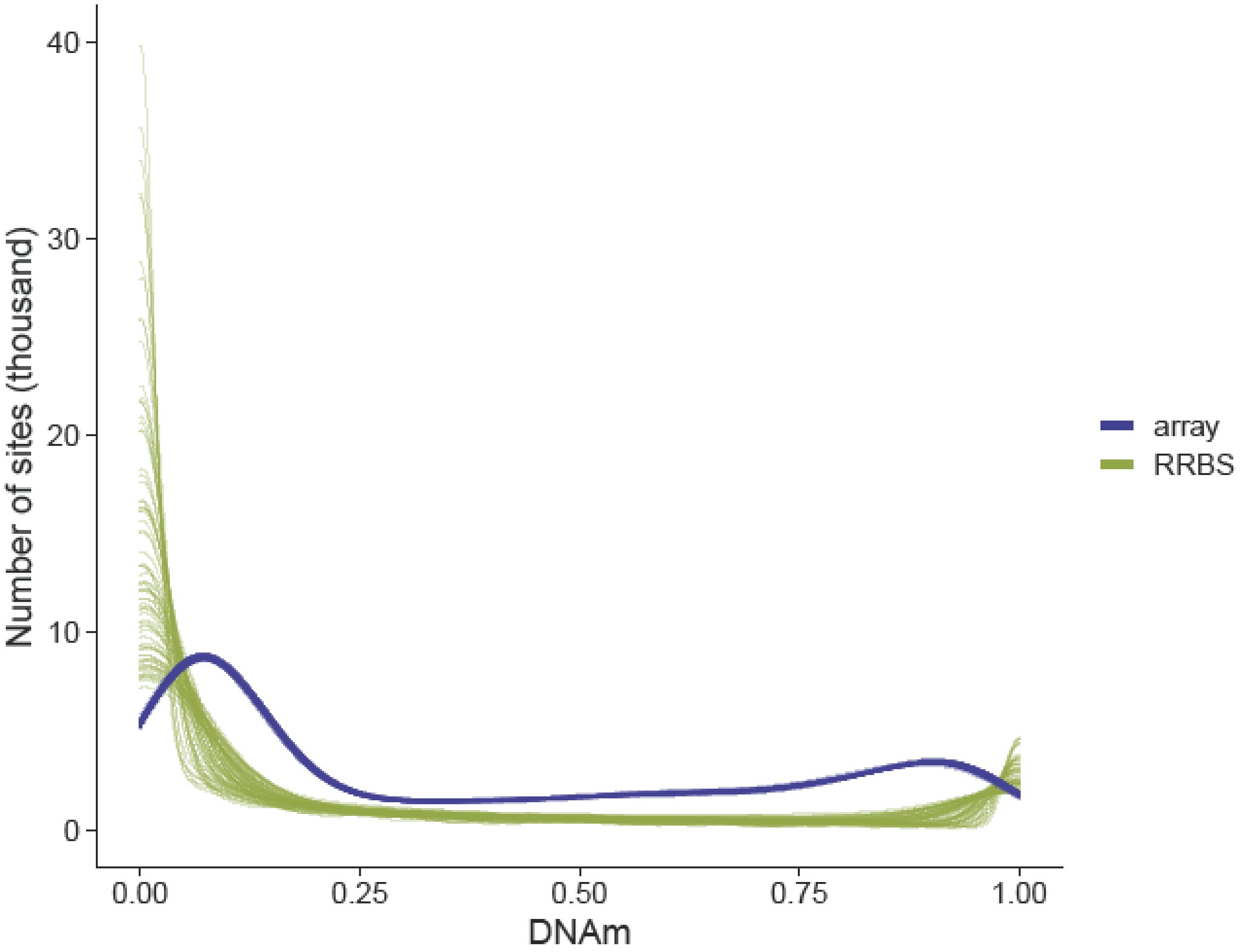

**Supplementary Figure 2: DNAm estimates derived from RRBS are on average lower than those from the array platform.** Shown are boxplots of the difference in DNAm estimated from the Illumina mammalian array and RRBS DNAm data across all overlapping DNAm points, with DNAm points grouped based on their read depths in the RRBS data.

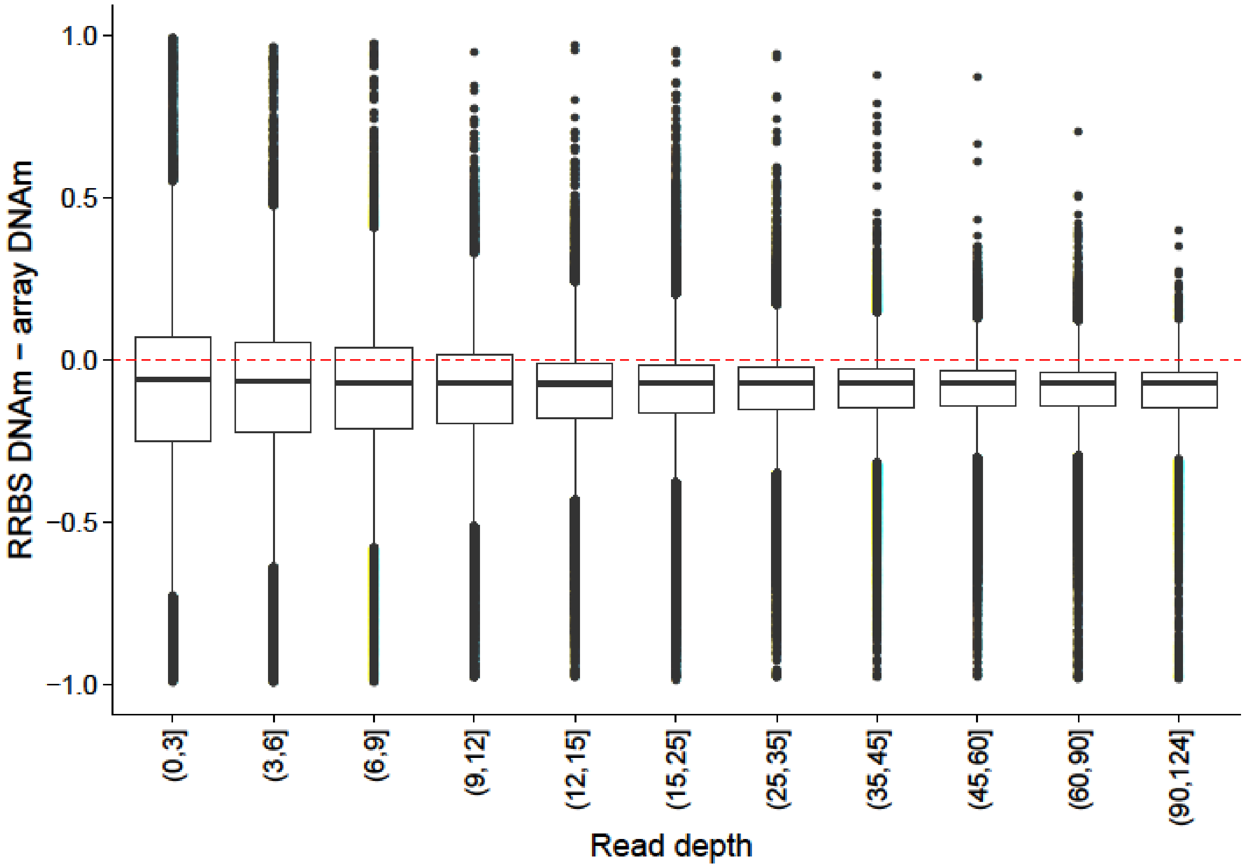

**Supplementary File 3: Outline of the framework for simulating bisulfite-sequencing data and assessing power in a DNAm site.** This framework can be expanded to simulate a range of different DNAm sites by varying the input parameters.

### WORKFLOW OF DNAM SITE-LEVEL POWER CALCULATION

*A power calculation for a two-group comparison for a DNAm site with characteristics matching the input parameters*

#### *INPUT*

| | |
|---|---|
| $\mathbf{N_1}$ | Sample size for group A |
| $\mathbf{N_2}$ | Sample size for group B |
| **µRD** | Mean read depth per group |
| **r** | Negative binomial function parameter |
| **µDNAm** | Mean DNAm |
| **ΔµDNAm** | Mean DNAm difference between groups |
| **nSites** | Number of DNAm sites of the same type to be simulated |
| **pValue** | P-value threshold applied to calculate power |

#### *1. Simulate DNAm sites*

i. Sample $\mathbf{N_1}$ + $\mathbf{N_2}$ read depths using the negative binomial distribution with mean read depth **µRD**, and parameter **r**. Resample any DNAm sites with a read depth of zero.

ii. Use the binomial distribution to sample DNAm level, where the probability is set to the mean of the group (group A = **µDNAm**, group B = **µDNAm ± ΔµDNAm**, bound between 0 – 1) and number of events is their read depth. Calculate the proportion of DNAm per by taking the mean of the binary values for each read.

iii. Repeat for **nSites** sites.

#### *2. Calculate power*

i. Use a two-sided t-test to compare the simulated DNAm of group A to that of group B.

ii. Power is the proportion of DNAm sites for which the t-test p-value is smaller than **pValue** .

**Supplementary File 4: Flow diagram describing the framework for simulating bisulfite sequencing studies utilized in POWEREDBiSeq.** An application of the framework described in **Supplementary Figure 3**, used to assess the power of a two-group bisulfite sequencing study given different filtering parameters.

### WORKFLOW OF STUDY-LEVEL POWER CALCULATION

*Simulating a two-group bisulfite sequencing study to calculate the power, given read depth and DNAm point filtering to be applied to the data, and expected DNAm difference*

| | |
|---|---|
| **RRBSTrue** | Unfiltered RRBS data matrix |
| **ΔμDNAm** | Expected mean difference in proportion of DNAm between groups |
| **nDNAmPoint** | Minimum number of DNAm points wanted in each group |
| **RDFilter** | The read depth filter to be applied |
| **pheno** | Binary variable indicating group membership (optional) |

#### *Inputs for DNAm site simulation*

| | |
|---|---|
| **$N_1$, $N_2$** | Equal to the number of samples in each group within **pheno**, or, if **pheno** is not given, as half of the total number of samples in **RRBSTrue**. |
| **μRD** | For the first **nDNAmPoint** DNAm points:<br>i. Calculate the mean read depth across **RRBSTrue**, using a subset of 60,000 DNAm sites.<br>ii. **μRD** is set to be the larger of **RDFilter** or mean read depth. Resample so that all have read depth > **RDFilter**.<br>For the remaining **$N_i$ – nDNAmPoint** DNAm sites:<br>i. **μRD** is the mean read depth, and sites with read depth < **RDFilter** to 0 to represent filtering. |
| **r** | Calculated using 60,000 sites and $\boldsymbol{r} = \frac{\frac{\mu^2}{\sigma^2}}{1-\frac{\mu}{\sigma^2}}$, where $\mu$ = mean read depth and $\sigma^2$ = variance of read depth across **RRBSTrue**. |
| **μDNAm** | i. Calculate the probability that DNAm is in ranges 0-0.05, 0.05-0.95, 0.95-1 using a subset of 100,000 DNAm sites from **RRBSTrue**.<br>ii. Sample the DNAm ranges, weighted by the probability calculated in (i).<br>iii. Use a uniform distribution to set **μDNAm** from values within the selected range. |
| **nSites** | One DNAm site of each type is simulated (**μDNAm** and **μRD** will differ for each DNAm site) |
| **pValue** | 0.05/(number of sites remaining in **RRBSTrue** after filtering by **RDFilter** and **nDNAmPoint**), a Bonferroni correction for the number of DNAm sites that would be compared. |

↓

#### *Use DNAm site simulation work flow to simulate a dataset*

i. Simulate 40,000 sites using the above inputs and step 1 of the workflow presented in **Supplementary Figure 3**.

ii. Bootstrap the p-values from the two-group t-test comparison so that you have the same number as the number of DNAm sites remaining in **RRBSTrue** after filtering by **RDFilter** and **nDNAmPoint**.

iii. Calculate study power using **pValue**.

**Supplementary Figure 5: A histogram of POWEREDBiSeq calculations showing variability in estimated power.** The estimated power of a study was calculated 400 times, using a mean DNAm difference between groups of 0.06 and minimum sample size of 60. Users can calculate bespoke power estimates using their own study-specific parameters in POWEREDBiSeq.

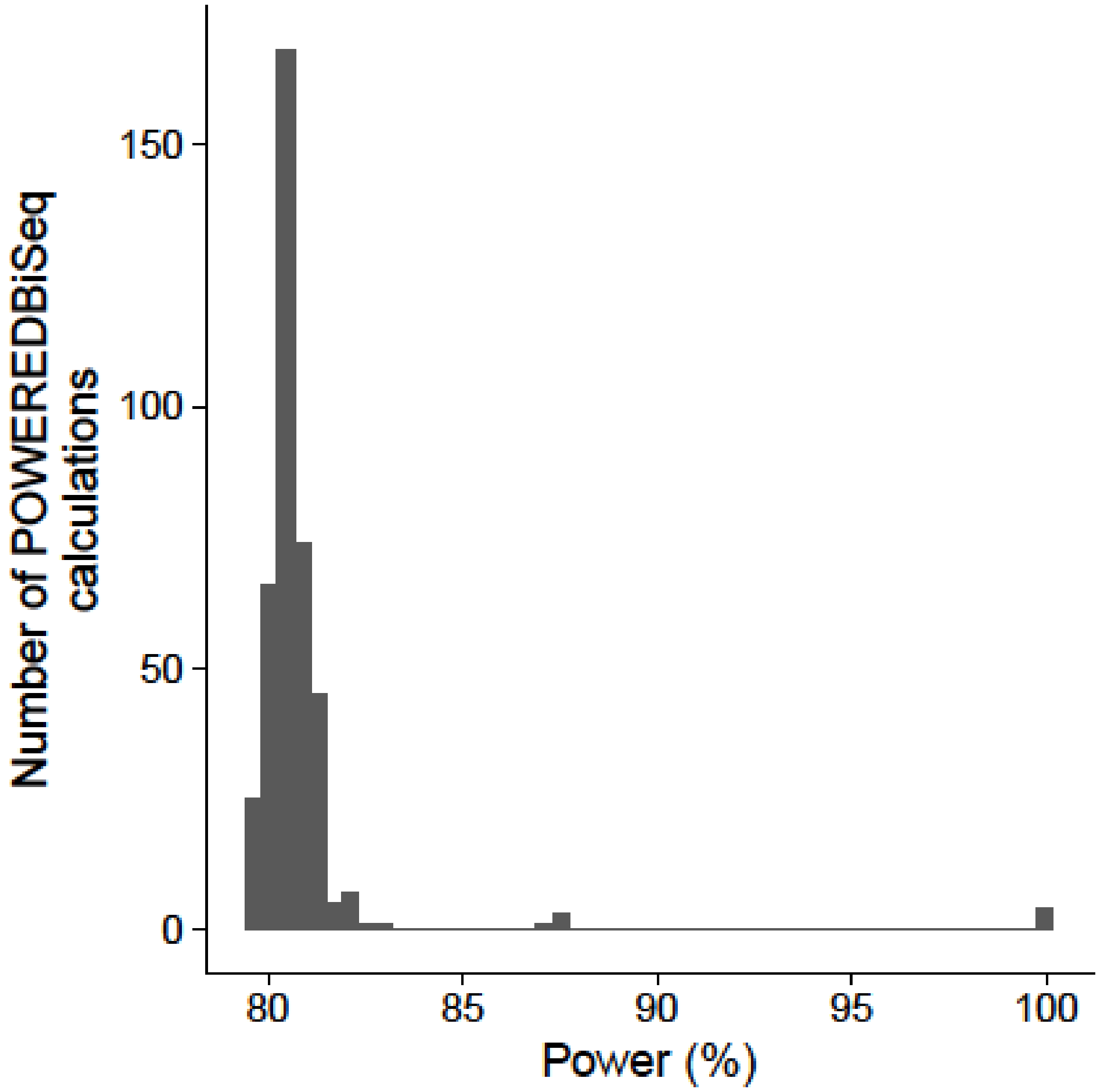

**Supplementary Figure 6: r is more accurately estimated when using a larger number of DNAm sites.** The negative binomial parameter, r, was calculated from an increasing number of DNAm sites using RRBS data from 125 samples. The red dashed line is the true value of r, calculated from the entire dataset.

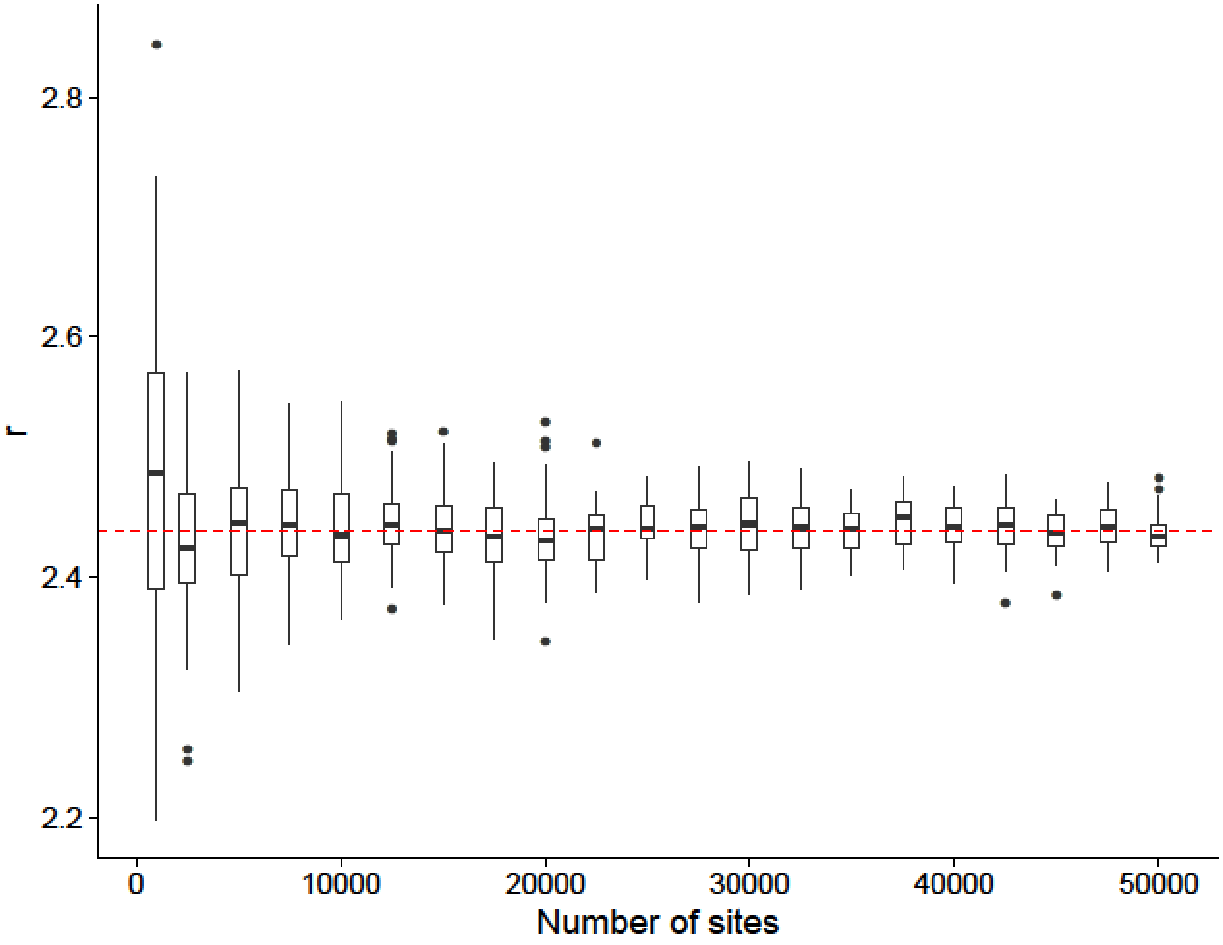

**Supplementary Figure 7: DNAm priors are more accurately estimated when using more DNAm sites.** DNAm priors are the probability that the DNAm value of selected DNAm sites fall within the DNAm bins 0-0.05, 0.05 - 0.95 and 0.95 – 1, shown in maroon, green and blue, respectively. Priors were calculated across 125 samples. The true value for each prior, calculated across the entire dataset, is shown in red.

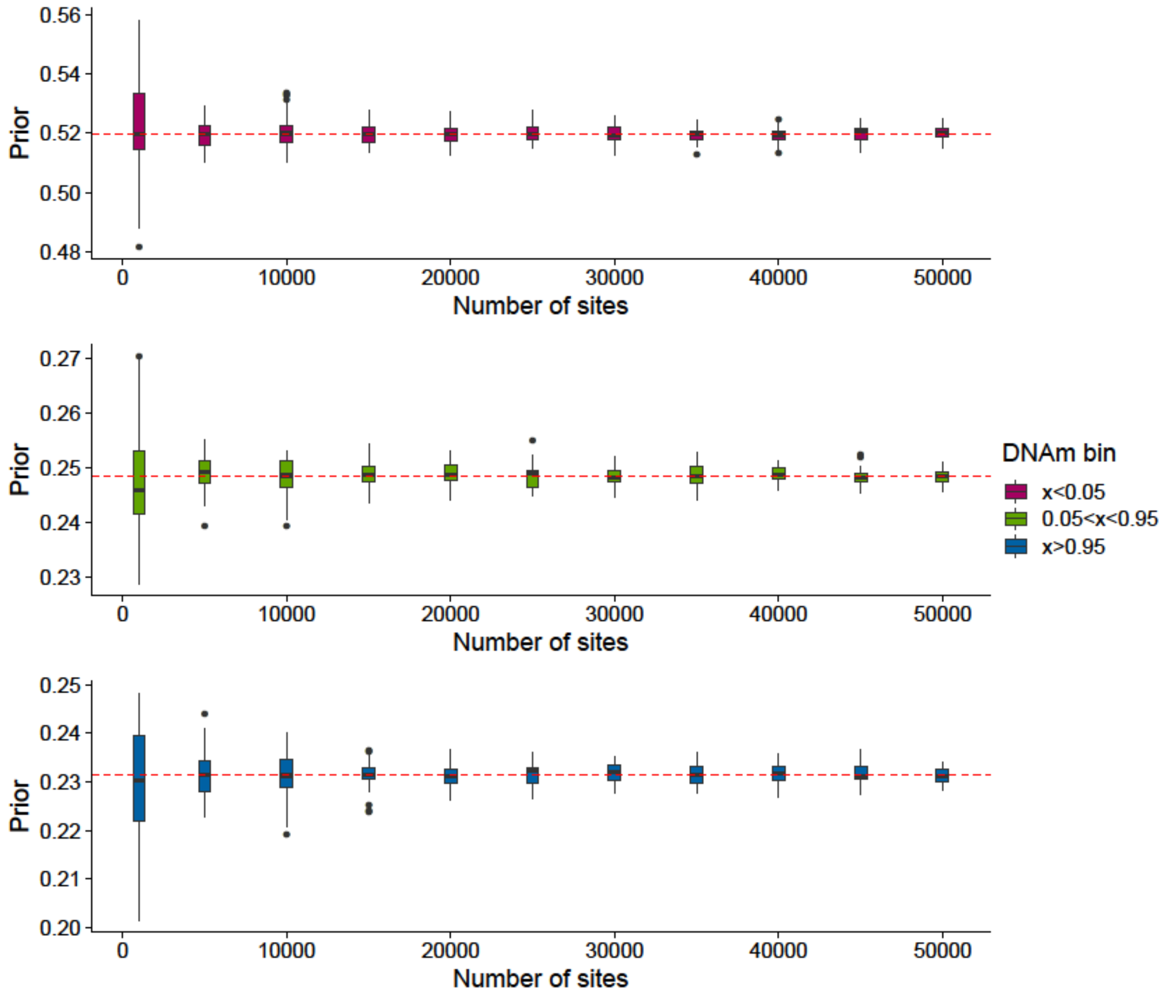

**Supplementary Figure 8: The proportion of DNAm sites remaining is more accurately estimated when using more DNAm sites.** The proportion of DNAm sites remaining after filtering by minimum read depth and minimum number of samples calculated from an increasing number of DNAm sites from an RRBS dataset. n was calculated across 125 samples. The red dashed line is the true value of n, calculated from the entire dataset.

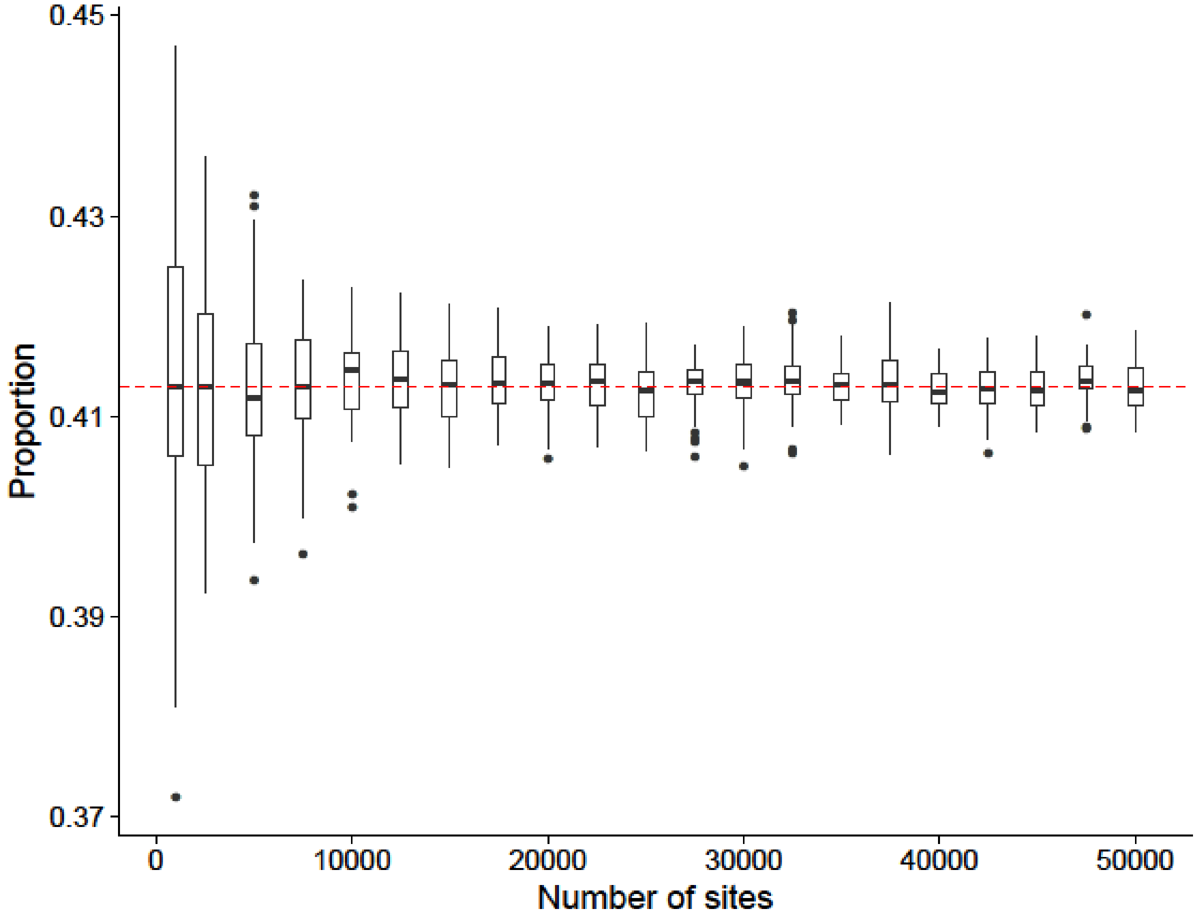

**Supplementary Table 1**: **A summary of parameters used in simulation analysis.**

| Plot | µRD | $N_1$, $N_2$ | ΔµDNAm | µDNAm |
|---|---|---|---|---|
| A | Variable | 30 | 0.2 | 0.25 |
| | | | 0.1 | |
| | | | 0.05 | |
| B | 20 | Variable | 0.2 | 0.25 |
| C | 25 | 20 | Variable | 0.25 |
| | | 10 | | |
| | | 5 | | |
| D, E | 50 | 80 | 0.05 | Variable |
| | 30 | | | |
| | 10 | | | |

Each row refers to a plot in **Figure 4**. Values were chosen so that the variable of interest could be seen to influence power within the scale of the figure.
